# Ergothioneine: Evaluation of a Novel Antioxidant for Targeting Ocular Oxidative Stress

**DOI:** 10.64898/2025.12.28.696576

**Authors:** Rujun He, Wei Ding, Juan Cao, Cong Guo, Xu Li, Guohua Xiao

## Abstract

**Purpose:** To evaluate L-Ergothioneine (EGT), a naturally occurring amino acid and endogenous antioxidant, as a novel therapeutic agent for oxidative stress-related ocular diseases. This evaluation specifically aimed to address the challenge of targeted ocular delivery by assessing EGT ‘s antioxidant potency, stability, ocular tolerance, and crucially, its ability to reach the posterior segment (fundus) via topical administration.

**Methods:** This study evaluated EGT as a novel ocular antioxidant by examining its radical scavenging capacity (DPPH assay compared to glutathione, astaxanthin, and coenzyme Q10), stability (at 40°C/75% relative humidity for six months using HPLC), ocular tolerance (using a New Zealand rabbit model), and fundus delivery efficiency (topical D_9_-EGT eye drops quantified by LC-MS/MS).

**Results:** EGT demonstrated significantly superior radical scavenging activity, exhibiting 6.4-fold and 46-fold higher rates than glutathione and coenzyme Q10, respectively, at 50 ppm. It also showed excellent stability, retaining over 97% of its initial concentration after six months, and caused no ocular irritation at any tested concentration (score 0). Importantly, topical administration of EGT resulted in effective fundus delivery, with peak concentrations reached at 0.5 hours post-application (1181 ± 56 ng/g), confirming successful penetration through corneal and scleral barriers. These findings establish EGT as a potent, multi-mechanistic antioxidant characterized by high stability, ocular safety, and exceptional posterior segment penetrance via non-invasive eye drops.

**Conclusion:** These findings establish EGT as a potent, multi-mechanistic antioxidant characterized by high stability, ocular safety, and exceptional posterior segment penetrance via non-invasive eye drops. By overcoming key delivery limitations, EGT presents a promising therapeutic strategy for oxidative stress-related ocular diseases such as age-related macular degeneration and diabetic retinopathy. Further studies are warranted to evaluate its long-term efficacy and clinical translation potential.

## 1. Introduction

Oxidative stress, defined as a state of damage caused by an imbalance between the production and clearance of reactive oxygen species (ROS) in the body, is a central pathological mechanism underlying the onset and progression of numerous ocular diseases[1]. Persistent oxidative damage can initiate or exacerbate several common blinding eye conditions, including age-related macular degeneration (AMD), cataracts [2], diabetic retinopathy (DR) [3], dry eye syndrome, and glaucoma [4]. Current clinical interventions, such as intravitreal anti-VEGF injections (for AMD and DR) [5], cataract surgery, and intraocular pressure-lowering medications (for glaucoma), can relieve symptoms or slow disease progression. However, these approaches primarily target middle-to-late disease stages and offer limited efficacy in addressing oxidative damage at its source during early stages [6]. Oral antioxidants (e.g., vitamins C and E, lutein, and zeaxanthin) are widely used [7], but their low ocular bioavailability and poor tissue targeting often prevent therapeutic concentrations from being achieved at the disease site [8]. Therefore, the development of efficient, low-toxicity antioxidants capable of penetrating ocular barriers and achieving targeted delivery remains a critical goal in ophthalmic drug research and development.

L-Ergothioneine (EGT), a naturally occurring sulfur-containing amino acid, has attracted considerable attention due to its unique antioxidant mechanisms [9]. EGT exhibits potent, multi-mechanistic antioxidant activity: it can directly and efficiently scavenge highly ROS such as hydroxyl radicals and hypochlorous acid [10]; chelate pro-oxidative divalent metal ions (e.g., Cu^2+^ and Fe^2+^) [11]; and indirectly enhance the expression of endogenous antioxidant enzymes, such as superoxide dismutase (SOD) and glutathione peroxidase (GSH-Px), by activating the Nrf2 signaling pathway, thereby forming a multi-layered antioxidant defense system [12]. In terms of targeting specific populations and alleviating symptoms, EGT can exert neuroprotective effects [13] and improve symptoms of diabetes and cardiovascular diseases [14] through functions such as antioxidant activity. Moreover, EGT exists as a zwitterion at physiological pH, and its oxidation products are relatively inert, capable of being reduced and regenerated, which prevents pro-oxidative effects.

The presence of a specific transporter in the human body, organic cation transporter novel type 1 (OCTN1), facilitates EGT uptake, contributing to its low cytotoxicity and excellent biocompatibility [15]. EGT accumulates in high concentrations in ocular tissues, including the lens, retina, cornea, and retinal pigment epithelium. Additionally, the gene encoding OCTN1 is highly expressed in various ocular tissues, such as the retina, cornea, and lens epithelium [16]. This widespread expression provides a molecular basis for the active uptake and retention of EGT in the eye, supporting its strong potential as a candidate for intraocular delivery.

## 2. Materials and Methods

### 2.1. Materials

EGT (purity >99%) was obtained from Jiangsu Gene III Biotechnology Co., Ltd. DPPH (CAS: 84077-81-6) was purchased from Sigma-Aldrich. Reduced glutathione (CAS: 70-18-8) and coenzyme Q10 (CAS: 303-98-0) were supplied by Shanghai Aladdin Biochemical Technology Co., Ltd. Astaxanthin oil (5.0% content) was provided by Yunnan Aierkang Biotechnology Co., Ltd. Ethanol (CAS: 64-17-5), high-performance liquid chromatography (HPLC)-grade acetonitrile (CAS: 75-05-8), sodium dodecyl sulfate (CAS: 151-21-3), and vitamin C (CAS: 50-81-7) were all purchased from Sinopharm Chemical Reagent Co., Ltd. New Zealand white rabbits were purchased from Vital River Laboratory Animal Technology Co., Ltd. and bred at Jiangsu Wanlue Pharmaceutical Technology Co., Ltd. D_9_-EGT (Cat. No. HY-N191451) was obtained from MedChemExpress. New Zealand white rabbits were purchased from Vital River Laboratory Animal Technology Co., Ltd. and bred at Jiangsu Wanlue Pharmaceutical Technology Co., Ltd. Detailed information of the rabbits was as follows: age ranged from 4 to 5 months, with the actual age deviation controlled within ±1 month at the time of administration; body weight ranged from 3 kg to 5 kg, with the actual weight deviation controlled within ±20% at the time of administration. These controls ensured the consistency of the animals’ physiological status and reduced experimental errors. All animal experiments were approved by the appropriate ethics committee and conducted in accordance with relevant guidelines (Approval No.: AP-202508).

### 2.2. Determination of antioxidant capacity

The antioxidant capacity in this study was assessed using the 1,1-diphenyl-2-trinitrophenylhydrazine (DPPH) assay, a widely used method known for its simplicity and visual color change (from purple to yellow). EGT and reduced glutathione were dissolved in deionized water, while astaxanthin and coenzyme Q10 were dissolved in ethanol. Each compound was prepared at a range of concentrations (10, 20, 30, 40, 50, and 60 mg/L). For the assay, 1 mL of each sample solution was mixed with either water or ethanol to reach a total volume of 3 mL. Then, 1 mL of DPPH ethanol solution (0.12 mg/mL) was added, and the mixture was thoroughly shaken and incubated in the dark at room temperature for 30 minutes. A mixture of pure water or ethanol with DPPH ethanol solution was used as the blank control. The absorbance was measured at 517 nm using a UV-Vis spectrophotometer, and the free radical scavenging capacity was calculated. Each concentration was tested in triplicate. The scavenging rate was calculated using the following formula [17]:

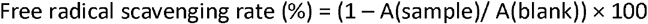

A (sample): Mixture of sample solution and DPPH solution;

A (blank): Mixture of pure solvent (without sample) and DPPH solution.

### 2.3. Stability experiment

An appropriate amount of EGT raw material was weighed, sealed in transparent polyethylene bags, and then placed into light-resistant composite aluminum bags (PET/Al/PE), with 50 g per bag. Simultaneously, a 1 mg/mL EGT solution was prepared using normal saline, sterilized by filtration, and dispensed into sterile glass bottles. Both the solid and solution samples were placed in a constant temperature and humidity chamber set at 40⍰± ⍰2°C and 75⍰± ⍰5% relative humidity for a duration of six months. Samples were collected at months 1, 2, 3, and 6. The EGT content was quantified using HPLC under the following chromatographic conditions: XDB-C18 column (250 × 4.6 mm, 5⍰μm), mobile phase of 5% acetonitrile in water, flow rate of 0.65 mL/min, detection time of 20 minutes, and a column temperature of 25°C. The retention rate was calculated based on the measured concentrations. Each sample and time point was analyzed in triplicate.

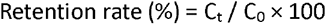

C_t_: Concentration at time t; C_0_: Initial concentration.

### 2.4. Acute ocular irritation test

New Zealand white rabbits (three per experimental group) were used as test subjects. Prior to the experiment, the eyes of all animals were examined 24 hours in advance. Rabbits exhibiting signs of ocular irritation, corneal defects, or conjunctival damage were excluded from the study. To administer the test substance, the lower eyelid of the rabbit’s left eye was gently pulled down, and 0.1 g or 0.1 mL of the test material was instilled into the conjunctival sac. The upper and lower eyelids were then passively closed for approximately 1 second to prevent loss of the test substance. The right eye remained untreated and served as a self-control. No rinsing was performed within 24 hours following administration. Clinical Observation: Ocular examinations were conducted at 1, 24, 48, and 72 hours after administration. If no irritation response was observed within 72 hours, the test was terminated. After the 24-hour observation, a 2% fluorescein sodium solution was applied to both eyes for further examination of corneal integrity. Ocular irritation was assessed according to standard eye damage scoring criteria, based on the mean scores and recovery times of the cornea, iris, and conjunctiva at 24, 48, and 72 hours post-application. A score of 0 for all three structures (cornea, iris, conjunctiva) was considered indicative of a non-irritating substance [18].

### 2.5. Rabbit eye fundus delivery test

Approximately 3 mg of D_9_-EGT was accurately weighed and dissolved in 0.6 mL of sterile normal saline for injection. After thorough mixing, a 5 mg/mL D_9_-EGT ophthalmic solution was obtained. New Zealand white rabbits were anesthetized, and their heads were gently tilted backward. The eyelids were carefully separated, and 50 µL of the eye drop solution was instilled into the right eye, or both eyes, using a pipette. At each time point, three rabbits were used, with five eyes receiving the D_9_-EGT eye drops and one eye serving as a control, receiving only sterile normal saline. During administration, the animals were maintained in a head-tilted position, and gentle pressure was applied to the nasolacrimal duct at the inner canthus for approximately 1 minute to prevent drainage of the solution through the nasolacrimal passage. The eyelids were intermittently closed to ensure full contact of the solution with the corneal and scleral surfaces. At 0.5 and 1 hour post-administration, ocular tissues, including aqueous humor, cornea, sclera, lens, vitreous body, and fundus, were collected. The concentration of D_9_-EGT in each tissue was quantified using liquid chromatography–tandem mass spectrometry (LC-MS/MS), enabling evaluation of its distribution and elimination across ocular compartments. All animal procedures were conducted in accordance with institutional ethical guidelines and were approved by the relevant ethics committee (Approval No.: AP-202508) [19]. All animal experiments were conducted in compliance with the standards set by AAALAC International accreditation and the ARRIVE guidelines. The experimental protocol was submitted to the Institutional Animal Care and Use Committee (IACUC) for review and approval prior to implementation. The use of animals adhered to the 3Rs principles (Reduction, Replacement, Refinement). All procedures related to experimental animals in this study were strictly performed in accordance with the international ethical code of conduct for laboratory animals, the “Guide for the Care and Use of Laboratory Animals. “The use of three animals per group was designed to minimize animal usage while ensuring the reliability of experimental results, which is consistent with standard practices for preliminary ocular pharmacology studies.

## 3. Results

### 3.1. Antioxidant capacity comparison test of EGT

As shown in Table 1, EGT demonstrated a markedly stronger free radical scavenging capacity compared to other tested antioxidants. At a concentration of 50 ppm, the antioxidant activity of EGT was 6.4 times higher than that of glutathione, 6.5 times higher than that of astaxanthin, and 46 times higher than that of coenzyme Q10.

**Table 1.** Comparison of free radical scavenging capacity between EGT and other antioxidants.

| Concentration<br>( PPM ) | DPPH Free Radical Scavenging (%) |  |  |  |
| --- | --- | --- | --- | --- |
|  | EGT | Glutathione (Reduced) | Astaxanthin | Coenzyme Q10 |
| 10 | 27.57±1.2 | 5.6±0.7 | 3.0±0.8 | 1.2±0.3 |
| 20 | 55.80±1.5 | 6.3±0.5 | 6.7±0.7 | 1.3±0.5 |
| 30 | 74.1±0.9 | 6.8±0.9 | 7.8±0.9 | 1.4±0.5 |
| 40 | 86.4±1.0 | 10.6±1.0 | 9.6±1.2 | 1.7±0.4 |
| 50 | 87.4±1.5 | 13.7±0.9 | 13.4±1.0 | 1.9±0.8 |
| 60 | 87.4±0.8 | 16.4±1.2 | 15.3±1.5 | 1.9±0.7 |

The DPPH free radical scavenging assay was used to determine the free radical scavenging rates (%) of ergothioneine (EGT), reduced glutathione, astaxanthin, and coenzyme Q10 at different concentrations (10–60 ppm). Data are presented as mean ± standard deviation.

### 3.2. Stability test of EGT

As shown in Table 2, the content of EGT in the raw material remained essentially unchanged after storage at 40°C for six months. Similarly, the retention rate of the EGT solution exceeded 97% under the same conditions, demonstrating its excellent stability and suitability for long-term storage.

**Table 2.** Stability results of EGT.

| Time<br>(Months) | Retention Rate of EGT Raw<br>Material<br>(%) | Retention Rate of EGT<br>Solution<br>(%) |
| --- | --- | --- |
| 1 | 99.9±0.2 | 99.9±0.1 |
| 2 | 100.0±0.2 | 99.0±0.1 |
| 3 | 100.0±0.1 | 98.5±0.3 |
| 4 | 100.0±0.0 | 98.1±0.1 |
| 5 | 100.0±0.2 | 97.6±0.2 |
| 6 | 100.0±0.0 | 97.2±0.4 |

A stability test was conducted under the conditions of 40°C and 75% relative humidity. High-performance liquid chromatography (HPLC) was used to determine the concentration retention rates (%) of ergothioneine (EGT) raw material and EGT normal saline solution at 1– 6 months. Data are presented as mean ± standard deviation.

### 3.3. Ocular irritation test of EGT

As shown in Table 3, EGT at various concentrations caused no ocular irritation in New Zealand white rabbits. Furthermore, no irritation was observed when EGT was incorporated into different eye care formulations, indicating excellent ocular tolerance across a range of applications.

**Table 3.** Results of acute ocular irritation responses of New Zealand white rabbits to EGT (three rabbits per group). The left eye of each rabbit received 0.1 g EGT pure powder or 0.1 mL 0.5% EGT eye wash via conjunctival sac instillation;

| Group | Eye Irritation Response Score |  |  |  | Results |
| --- | --- | --- | --- | --- | --- |
|  | 1h | 24h | 48h | 72h |  |
| Control | 0 | 0 | 0 | 0 | Non-irritating |
| EGT Pure Powder | 0 | 0 | 0 | 0 | Non-irritating |
| 0.5% EGT Eye Wash | 0 | 0 | 0 | 0 | Non-irritating |

### 3.4. Rabbit eye delivery test of EGT

As shown in Table 4, after New Zealand white rabbits were topically administered 5 mg/mL D_9_-EGT eye drops, liquid chromatography-tandem mass spectrometry (LC-MS/MS) detection revealed that the concentration of D_9_-EGT in various ocular tissues peaked at 0.5 hours post-administration and decreased at 1 hour, with the fundus maintaining a detectable therapeutic concentration at both time points. Specifically, the concentrations of D_9_-EGT in the cornea were 3060 ± 76 ng/g and 2700 ± 58 ng/g at 0.5 hours and 1 hour, accounting for 0.0904 ± 0.0333% and 0.0734 ± 0.0391% of the total instilled amount, respectively, which were at high levels among all tissues. The concentrations in the sclera were 1181 ± 56 ng/g and 217 ± 31 ng/g at 0.5 hours and 1 hour, accounting for 0.2450 ± 0.0581% and 0.0355 ± 0.0144% of the total instilled amount. The concentrations in the fundus were 589 ± 36 ng/g and 306 ± 23 ng/g at 0.5 hours and 1 hour, accounting for 0.0208 ± 0.0169% and 0.0091 ± 0.0057% of the total instilled amount. D_9_-EGT was also detected in the aqueous humor and lens, but not in the vitreous humor. These results clearly confirm that EGT can be effectively delivered to the posterior segment tissues of the eye including the fundus via topical eye drop administration.

**Table 4.**
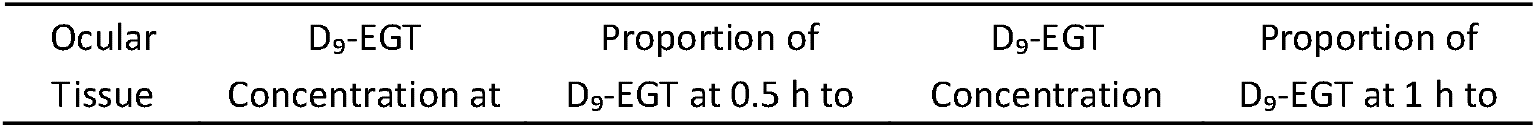
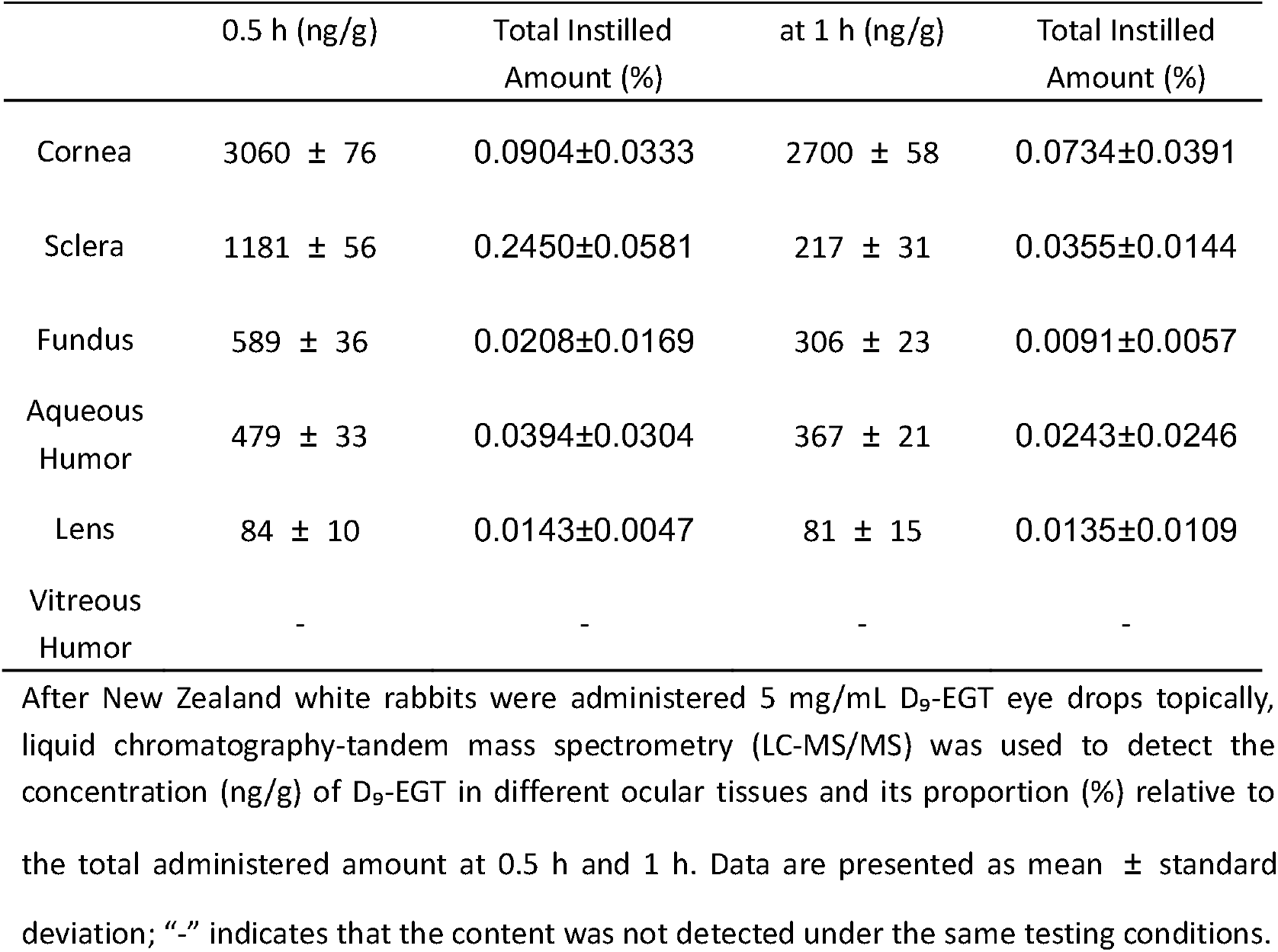
Distribution of D_9_-EGT in various parts of rabbit eyes at different time points after topical administration.

## 4. Discussion

Current drug delivery strategies for oxidative stress-related fundus diseases, such as AMD and DR, face significant challenges. Oral administration is severely limited by the blood–retinal barrier (BRB), resulting in minimal drug penetration into intraocular tissues [20]. Although intravitreal injection enables direct delivery to the posterior segment, it is an invasive procedure associated with potential complications, including infection, intraocular hemorrhage, and retinal detachment. Additionally, the short intraocular half-life of many drugs necessitates frequent injections, which negatively impacts patient compliance. Topical eye drops are the most convenient and non-invasive method of ocular drug delivery. However, conventional small-molecule drugs and biologic macromolecules (e.g., antibodies) exhibit extremely low delivery efficiency to the fundus, typically less than 0.01%, due to multiple barriers, including the cornea, tear drainage, and the BRB [21]. This severely limits their therapeutic efficacy for posterior segment diseases.

This study demonstrates that EGT offers unique advantages in the treatment of oxidative stress-related ocular diseases. Its potent ROS scavenging ability, metal ion chelation capacity, and activation of endogenous antioxidant defense systems provide a comprehensive strategy for combating complex oxidative damage in the eye. Thermal stability experiments confirmed the excellent storage performance of EGT formulations, while acute ocular irritation tests validated their high local safety and non-irritating properties, supporting suitability for long-term use. Crucially, ocular delivery studies in rabbits confirmed that topical administration of EGT eye drops can overcome multiple anatomical barriers and achieve effective delivery to the fundus. This enhanced delivery is largely attributed to the high expression of the OCTN1 transporter in ocular tissues, particularly in the retina, enabling active, targeted transport and overcoming the delivery efficiency limitations of conventional eye drops. As a naturally occurring compound that is readily absorbed and utilized by the human body, EGT also offers superior biocompatibility compared to many synthetic antioxidants, further reinforcing its potential as a safe and effective therapeutic candidate.

Notably, D_9_-EGT was undetectable in the vitreous humor of rabbits following topical administration, which may be explained by two key factors. First, topically delivered drugs (e.g., eye drops) require sequential penetration of ocular barriers: via the corneal pathway (crossing epithelial and stromal layers) or conjunctival-scleral pathway (penetrating conjunctival epithelium and sclera) to reach the posterior segment. The retina, a choroid-attached transparent tissue anterior to the vitreous humor, acts as a spatial barrier —drugs must first penetrate retinal tissues before diffusing into the vitreous humor behind, delaying or restricting distribution[22]. Second, the vitreous humor, a high-viscosity gelatinous matrix composed of water, collagen, and hyaluronic acid, inherently impedes small-molecule diffusion[23]. Although EGT (≈ 229 Daltons) exhibits favorable corneal permeability, it exists as a zwitterion at physiological pH (isoelectric point ≈5.5), reducing lipid solubility and limiting passive diffusion across the lipid-rich BRB, thereby further hindering vitreous entry[24].

Based on these advantages, EGT shows broad application potential in the treatment and prevention of ocular diseases associated with oxidative stress. As a topical eye drop, it may be used in high-risk populations, such as the elderly and individuals with diabetes, to prevent the onset and progression of cataracts, AMD, and DR. It also holds promise as an adjunctive therapy alongside existing treatments, such as anti-VEGF agents, where it may synergistically enhance therapeutic outcomes by mitigating oxidative damage and reducing vascular leakage in AMD and DR. Due to its potent antioxidant and anti-inflammatory properties, along with favorable corneal permeability, EGT is also a promising candidate for managing dry eye syndrome, where it may alleviate oxidative damage and inflammatory responses. Furthermore, its neuroprotective potential offers a novel strategy for glaucoma by protecting retinal ganglion cells from oxidative stress-induced degeneration.

Incorporating EGT into advanced drug delivery systems, such as liposomes, nanoparticles, or sustained-release gels, could further prolong ocular residence time and increase drug concentrations in target tissues, thereby optimizing therapeutic efficacy.

## 5. Conclusions

This study systematically evaluated the potential of EGT as an ocular antioxidant therapy. DPPH assays and stability tests confirmed its strong free radical scavenging capacity and excellent chemical stability. Acute ocular irritation assessments and fundus delivery experiments demonstrated that EGT eye drops are safe, non-irritating, and, crucially, capable of overcoming ocular barriers to reach target tissues such as the fundus following topical administration. Compared with the limitations of current therapeutic agents, EGT offers significant advantages, including efficient and multi-mechanistic antioxidant activity, excellent ocular tissue permeability (notably via OCTN1-mediated active transport to the fundus), strong stability, and high ocular safety. Collectively, these findings position EGT as a highly promising candidate for the prevention and treatment of a range of blinding ocular diseases related to oxidative stress, including AMD, DR, cataracts, and dry eye syndrome.

The present study has a few limitations. First, while untreated eyes (self-controls) and normal saline-treated eyes (negative controls) validated EGT’s ocular safety and delivery efficiency, standard clinical antioxidant eye drops (e.g., those containing vitamin C, lutein, or astaxanthin) were not included as positive controls. This omission precludes direct comparison of EGT’s therapeutic potential with clinically established antioxidants; Second, only two post-administration time points (0.5 h, 1 h) were analyzed for D_9_-EGT, leaving key pharmacokinetic parameters [elimination half-life (t_1_/_2_), area under the concentration-time curve (AUC), fundus drug residence time] undefined and hindering dosing optimization; Third, and although EGT reached the fundus (0.5 h: 589 ± 36 ng/g; 1 h: 306 ± 23 ng/g), the therapeutic relevance of these concentrations remains unvalidated—specifically, whether they suffice for antioxidant, anti-inflammatory, or neuroprotective effects, and how long such effects persist. This gap weakens direct evidence for EGT’s clinical application, requiring follow-up studies with oxidative stress-related ocular disease models to establish concentration-efficacy correlations.

Future studies should focus on verifying its long-term efficacy in relevant ocular disease models, optimizing delivery formulations, and advancing toward clinical translation. Additionally, further research could explore the therapeutic potential of EGT in comorbidity models of ocular diseases and neurodegenerative disorders, leveraging its established value in cognitive protection to provide new strategies for the combined treatment of cross-system oxidative stress-related diseases.

## Availability of data

The data that support the findings of this study are available from the corresponding author, upon reasonable request.

## Conflict of Interest

The authors declare no competing financial interests.

## Ethical Statement

All animal experiments were approved in accordance with ethical guidelines (No.: AP-202508).

## Acknowledgements

We would like to acknowledge Gene III Biotechnology Co., Ltd. for providing the research sponsor. We also acknowledge Jiangsu Wanlue Pharmaceutical Technology Co., Ltd. for resource docking in fundus delivery tests.

---

**Figure 1.**
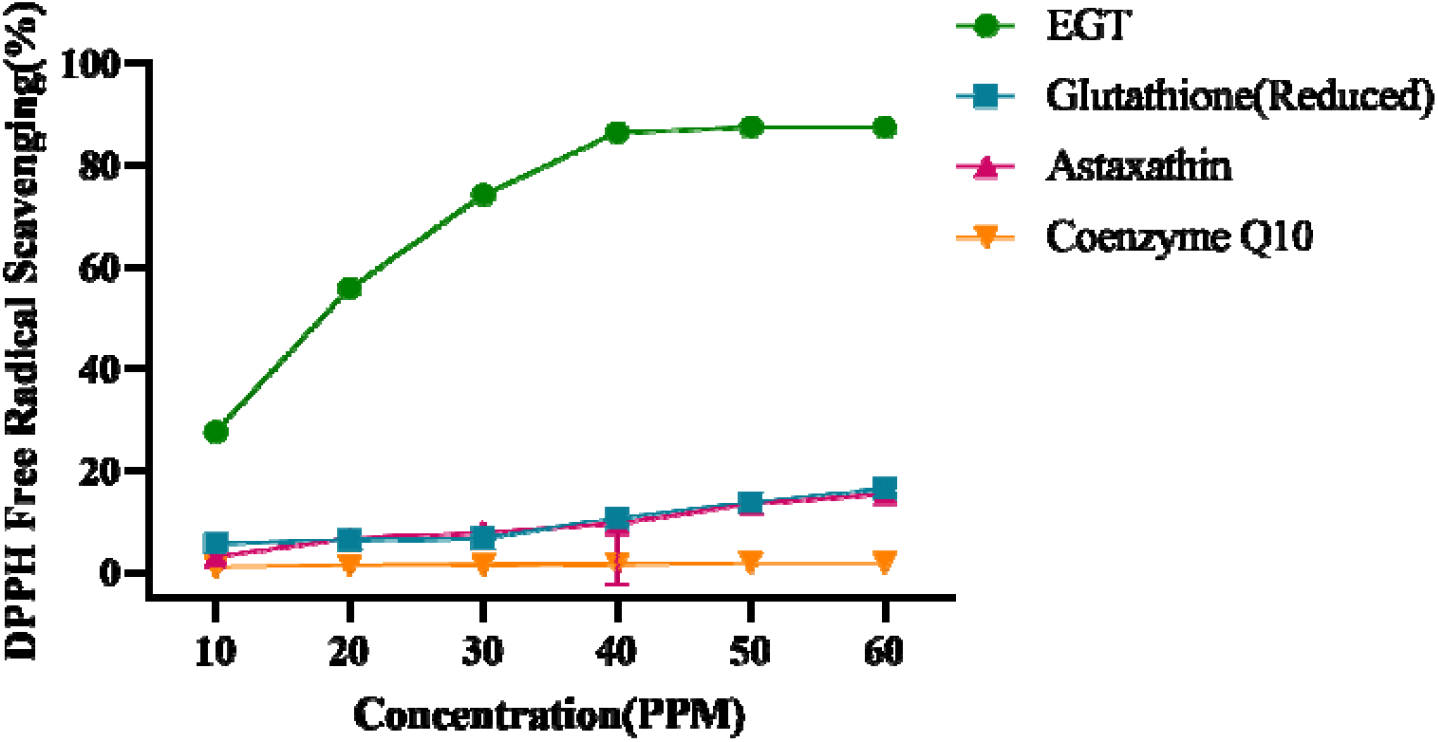
Comparison of DPPH free radical scavenging rates of ergothioneine (EGT) and three other common antioxidants (reduced glutathione, astaxanthin, and coenzyme Q10) at concentrations ranging from 10 to 60 ppm.

**Figure 2.**
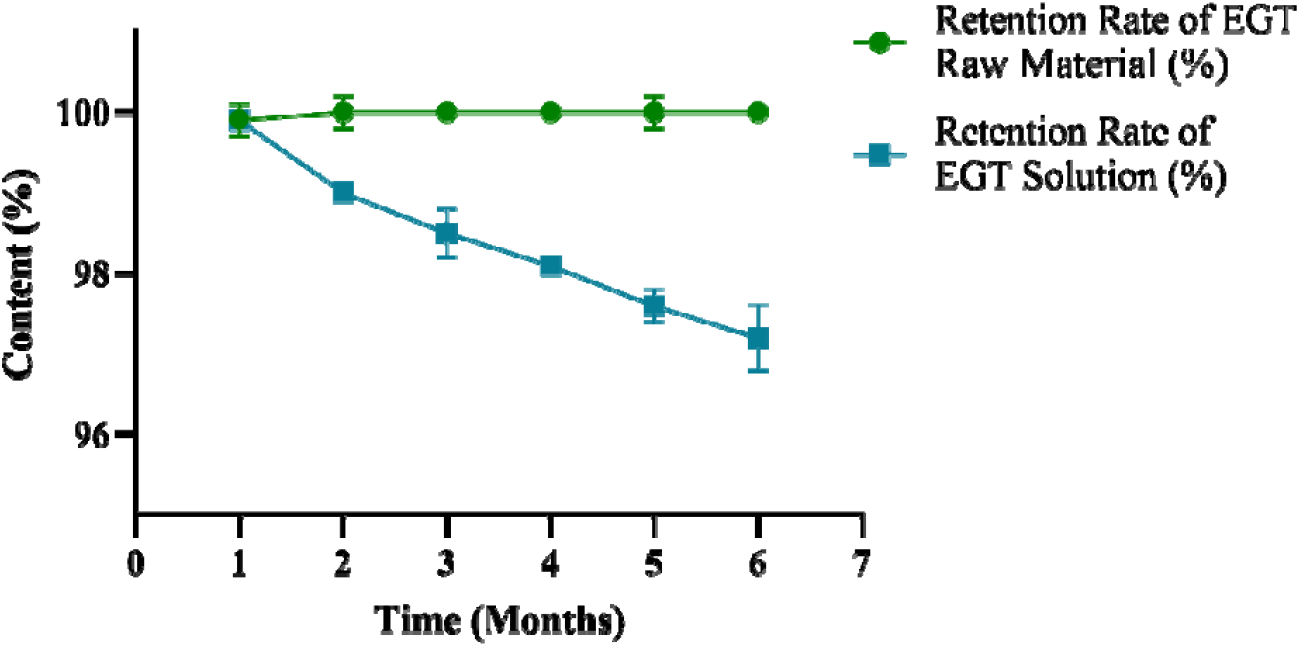
Results of retention rates of EGT in raw material and solution forms over 6 months of storage under accelerated conditions (40 ± 2°C, 75 ± 5% relative humidity).

**Figure 4.**
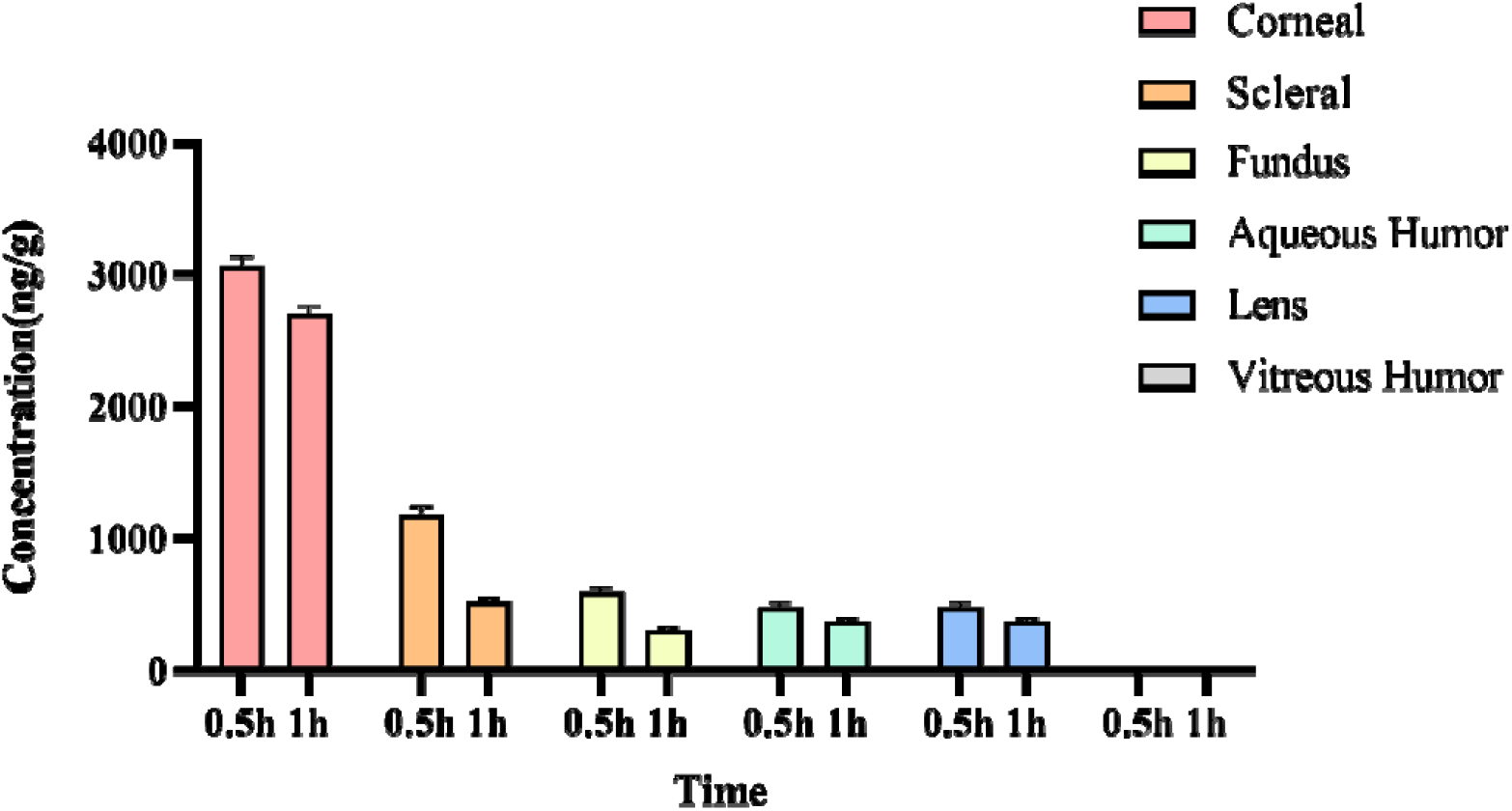
the concentrations of deuterium-labeled EGT (D_9_-EGT) in different rabbit ocular tissues at 0.5 and 1 hour post-topical administration.

## Notes

### Competing Interest Statement

The authors have declared no competing interest.

